# Chronic Dietary Exposure to Methylparaben and Ethyl paraben Induces Developmental, Biochemical, and Behavioural Toxicity in Drosophila melanogaster

**DOI:** 10.1101/2025.10.14.682250

**Authors:** Ravish Huchegowda, Shreesha Simha Bhat, Prathusha Srinivas, Meghana Tare, Dhruti Reddy Pradeep, S R Sahana, Rahul Dubey, RK Ranganath, P R Manjunath

## Abstract

Parabens, particularly methylparaben (MP) and ethylparaben (EP), are extensively used preservatives in cosmetics, foods, and pharmaceuticals. Although considered safe at low concentrations, recent evidence questions their biological inertness under chronic exposure. This study evaluated the developmental, biochemical, and behavioral effects of continuous dietary MP and EP exposure in *Drosophila melanogaster*, an established in vivo model for toxicological screening. Flies were chronically exposed to MP (0.5–2%) or EP (0.5–1.5%) throughout development and adulthood. Developmental timing, lifespan, oxidative-stress markers (MDA, FRAP, total protein), and locomotor performance (negative geotaxis in adults, crawling in larvae) were quantified. Paraben exposure significantly delayed development (∼15% increase in eclosion time), reduced median lifespan (up to 50% decrease at 2% MP), and elevated oxidative damage (↑MDA, ↓FRAP) in a dose-dependent manner. Protein content declined more rapidly with age, suggesting oxidative degradation or proteolysis. Both adult climbing and larval crawling performances were impaired, linking biochemical stress to neuromuscular dysfunction. MP produced stronger oxidative and behavioral effects than EP. Feeding controls confirmed that observed deficits were not due to nutritional differences. Chronic MP and EP exposure induces systemic toxicity in *D. melanogaster*, integrating endocrine disruption and redox imbalance as plausible mechanisms. Given conserved stress and hormonal pathways, these findings reinforce the need to re-evaluate low-dose paraben safety limits and highlight *Drosophila* as a rapid, ethically viable platform for screening environmental preservatives and safer substitutes.

## 1. Introduction

Parabens especially methylparaben (MP) and ethylparaben (EP) are ubiquitous preservatives in cosmetics, processed foods, and pharmaceuticals. Human exposure is continuous via dermal, oral, and inhalational routes, and parabens are detectable in urine, blood, breast milk, placenta, and other tissues(Haldar et al., 2025) (Hager et al., 2022). Although historically permitted at low levels (≤0.4% for any single paraben and ≤0.8% in combination in cosmetics), accumulating evidence indicates that chronic low-dose exposure may not be biologically inert(Torfs & Brackman, 2021; Wei et al., 2021). Mechanistically, parabens act as endocrine-disrupting chemicals: they exhibit weak estrogenicity through ERα/ERβ binding, perturb the hypothalamic– pituitary–thyroid axis, and display anti-androgenic activity; they also elevate reactive oxygen species, promoting lipid peroxidation and protein oxidation, with neurobehavioral consequences reported in vertebrate models (Azeredo et al., 2023; Kim et al., 2024; Kley et al., 2024; Nagar et al., 2020). *Drosophila melanogaster* offers a tractable in vivo system to interrogate these risks due to conserved disease-gene orthologs, rapid life cycle, and validated assays for development, stress biology, and behavior; Most studies focus only on developmental or molecular outcomes, making it hard to connect mechanisms across biological levels. We chronically expose flies to dietary MP or EP and measure development, lifespan, oxidative-stress markers (total protein, lipid peroxidation, antioxidant capacity), and locomotor performance. This study aimed to evaluate whether chronic dietary exposure to methyl- and ethyl-paraben induces measurable toxicity across developmental, biochemical, and behavioral domains in *Drosophila melanogaster*.

## Materials and Methods

### 2.1. Experimental design

We assessed the effects of chronic *dietary methylparaben* (MP) or ethylparaben (EP) exposure in *Drosophila melanogaster* by measuring development time, adult lifespan, markers of oxidative stress and protein levels, and locomotor abilities. All assays were conducted under consistent environmental conditions unless specified. The number of replicates per group (n = 5 vials) was based on prior Drosophila toxicology studies showing that this replication level achieves >80 % power for detecting ≥15–20 % changes in survival or biochemical markers at α = 0.05. Thus, the sample size used here was adequate to detect biologically meaningful effects while minimizing animal use.

### 2.2. Fly stocks and husbandry

Wild-type *D. melanogaster* (Canton-S) were maintained on standard cornmeal–yeast–agar food in ventilated vials at 18 °C, 50–60% RH, 12:12 h light:dark. Adults were density-matched (12 flies per vial; 1:1 sex ratio post-eclosion unless otherwise specified).

### 2.3. Diet preparation and paraben dosing

MP and EP stocks were prepared fresh in ethanol and added to cooled molten food to yield final concentrations (w/v): MP: 0.5%, 1.0%, 2.0%; EP: 0.5%, 1.0%, 1.5%. Final ethanol did not exceed 1% v/v and was matched in all groups. All media including control and paraben-supplemented—contained TEGO 1000 mg/L (same lot and concentration) to keep the antimicrobial background identical. Treatment concentrations for MP/EP were in addition to this base.

### 2.4. Cohorting and exposure paradigm

For oviposition, 12 adults were introduced into each vial (control or paraben food) for 48 h and then removed to ensure continuous exposure from embryo to adult. After eclosion, 12 newly eclosed adults were maintained per vial on their respective diets and transferred to fresh food twice weekly.

### 2.5. Developmental timing

Larval development was monitored daily. Time to pupation and time to adult eclosion were recorded for each group (using n>100 individuals per condition) as measures of developmental progression. To assess adult longevity, cohorts of 50 newly eclosed flies per group (mixed sex, equal ratio) were transferred to fresh treatment or control food and monitored for survival. Deaths were scored daily, and surviving flies were transferred to fresh food twice weekly. Survival data were used to generate Kaplan–Meier survival curves, and median lifespan was calculated for each group.

### 2.6. Biochemical Measurements

We collected fly heads at early-adult (∼5 days), mid-life (∼20 days), and late-life (∼30 days) stages (10 heads per sample, 5 samples per group). Heads were chosen for biochemical assays to focus on neuromuscular/brain function and reduce interference from gut content, pigments, and reproductive tissues, aligning with behavioral measures. Total protein was determined by Bradford assay(Bradford, 1976), lipid peroxidation by TBARS (MDA equivalents)(Ohkawa et al., 1979), and antioxidant capacity by FRAP (μmol Fe^2+ equivalents/mg protein)(Benzie & Strain, 1996). All spectrophotometric readings were performed in triplicate.

### 2.7. Behavioral Assays

#### 2.7.1. Adult locomotor

abilities were measured using the negative geotaxis assay, which involves startle-induced climbing. Groups of 12 adult flies from each treatment and control group were tapped to the bottom of a vertical tube, and their climbing activity was video-recorded. The footage was analyzed with Ctrax software to determine average climbing speed and performance(Branson et al., 2009). Each group underwent five separate replicates, and average results were used to generate graphs in Graph Pad Prism for statistical analysis.

#### 2.7.2. Larval movement

was assessed with a crawling assay in late third-instar larvae. Each larva was placed on a 1% agar plate, and its movement was recorded for 1 minute. Videos were evaluated using EthoVision software to measure average crawling speed and total distance travel(Spink et al., 2001). Both treatment and control groups had five replicates, and data visualization and statistical tests were carried out in Graph Pad Prism. All larval velocities are reported in mm/s to avoid ambiguity (0.1–0.2 cm/s = 1–2 mm/s). Calibration used the plate’s 10 mm scale; any trials with tracking loss or edge contact >10% of frames were excluded a priori.

To ensure that changes in behavior or biochemistry were linked to paraben exposure rather than reduced feeding or starvation, a feeding verification assay was conducted. This involved mixing a non-toxic food dye into the food given to separate cohorts of both control and paraben-treated larvae and adults for 24 hours. Visual inspection confirmed dye presence in the digestive tracts of all groups, verifying that feeding was unaffected. Thus, the observed behavioral and biochemical effects could be attributed specifically to paraben exposure.

### 2.8. Feeding verification

To rule out starvation, red food-grade dye was added to control and treatment media; larval midguts and adult crops were examined after 24 h exposure. Images are presented in **Supplementary Figure S1**.

### 2.9. Reagents and instrumentation

MP and EP (analytical grade); TEGO (antimicrobial), ethanol (molecular biology grade), BSA (Bradford), TBA and MDA standards (TBARS), FRAP reagents (ferric chloride, TPTZ). CO□-free handling for behavior; dissecting microscope; microplate reader (absorbance for Bradford/TBARS/FRAP); video cameras ≥30 fps; tracking software (Ctrax, EthoVision XT).

## 3. Data Analysis

All quantitative data are presented as mean ± standard error. Developmental timing and behavioral endpoints were compared between groups by one-way ANOVA or Kruskal–Wallis test (for non-normal distributions), with Dunnett’s or Dunn’s post hoc tests to compare each treatment to control. Survival curves were compared using the log-rank test (Mantel–Cox). Biochemical assay results were analyzed by ANOVA with Tukey’s multiple comparisons. A threshold of *p*<0.05 was considered statistically significant for all tests. Statistical analyses were performed using GraphPad Prism 9.

### 3.1. Randomization, blinding, exclusion criteria, and replication: Randomization

Vials were grouped using a random number generator; assay order was randomized daily. Random sequence generation was performed in Microsoft Excel (RAND function) before the start of exposure. Each vial (n = 5 per group) was assigned a unique random number, which determined its allocation to control or paraben-supplemented food (MP 0.5 %, 1 %, 2 %; EP 0.5 %, 1 %, 1.5 %).

Investigators performing behavioral analysis and biochemical quantification remained blind to group identity until data processing was complete. Adults and larvae were transferred to coded vials labeled only with random IDs, ensuring allocation concealment during subsequent handling and blinding during analysis. The same randomization file was used to randomize assay order (behavioral and biochemical measurements) across days to minimize temporal bias. **Blinding:** Video files were coded so scorers were blind to treatment during tracking and QC. **Exclusions (pre-registered):** Excluded vials had contamination or >20% mortality within 48 h post-eclosion; tracking runs with glare or >10% frame loss; larvae contacting the plate edge in >10% of frames. **Replication:** Unless specified, each group per assay had ≥5 biological replicates (vial as unit) and technical triplicates for spectrophotometric reads.

## 4. Results

### 4.1. Paraben Exposure Delays Development and Reduces Adult Survival

Chronic exposure to MP or EP significantly delayed *D. melanogaster* development, with higher doses (2% MP, 1.5% EP) extending time to eclosion by ∼2 days (∼15%, *p*<0.01). Lower concentrations also caused significant, though smaller, developmental delays. Paraben-treated groups exhibited reduced median and maximum lifespans in a dose-dependent manner; 2% MP decreased mean lifespan by 50% (median: ∼20 vs. ∼40 days in controls), while 1.5% EP shortened lifespan by ∼30%. Survival curves differed significantly from controls at all d*o*ses (p<0.001 for *h*igh, p<0.01 for low). These findings show that chronic paraben exposure disrupts development and reduces longevi*ty in Drosophila Melanogaster*. Effect-size estimation and confidence intervals For each primary and secondary outcome, mean or median differences between treatment and control groups were computed along with 95 % confidence intervals using GraphPad Prism 9. Where applicable, effect sizes (Cohen’s *d*, η^2^, or hazard ratios for survival) were also calculated to express the magnitude of paraben-induced change. Reporting both p-values and CIs allows direct assessment of the precision and biological relevance of each effect. **(Figure 2a and 2b) (Table 1)**.

**Table 1.** Developmental Progress in *D. melanogaster* Under Methyl paraben (MP) and Ethyl paraben (EP) Exposure.

| Group | Avg. Time to Pupation (Days) | Avg. Time to Eclosion (Days) | Developmental Delay | Remarks |
| --- | --- | --- | --- | --- |
| Control | 10–11 | 18–20 | None | Normal pupation and adult emergence |
| 0.5% MP | 11–12 | 20–21 | Mild | Slight delay in both stages |
| 1.0% MP | 12–13 | 22–23 | Moderate | Delayed pupation and fewer eclosed adults |
| 2.0% MP | 13–14 | 24–25 | Severe | High larval retention, very few eclosed |
| 0.5% EP | 11–12 | 20–21 | Mild | Similar pattern as 0.5% MP |
| 1.0% EP | 12–13 | 22–23 | Moderate | Reduced pupation and delayed emergence |
| 1.5% EP | 13–14 | 24–25 | Severe | Developmental arrest in many larvae |

**Figure 1.**
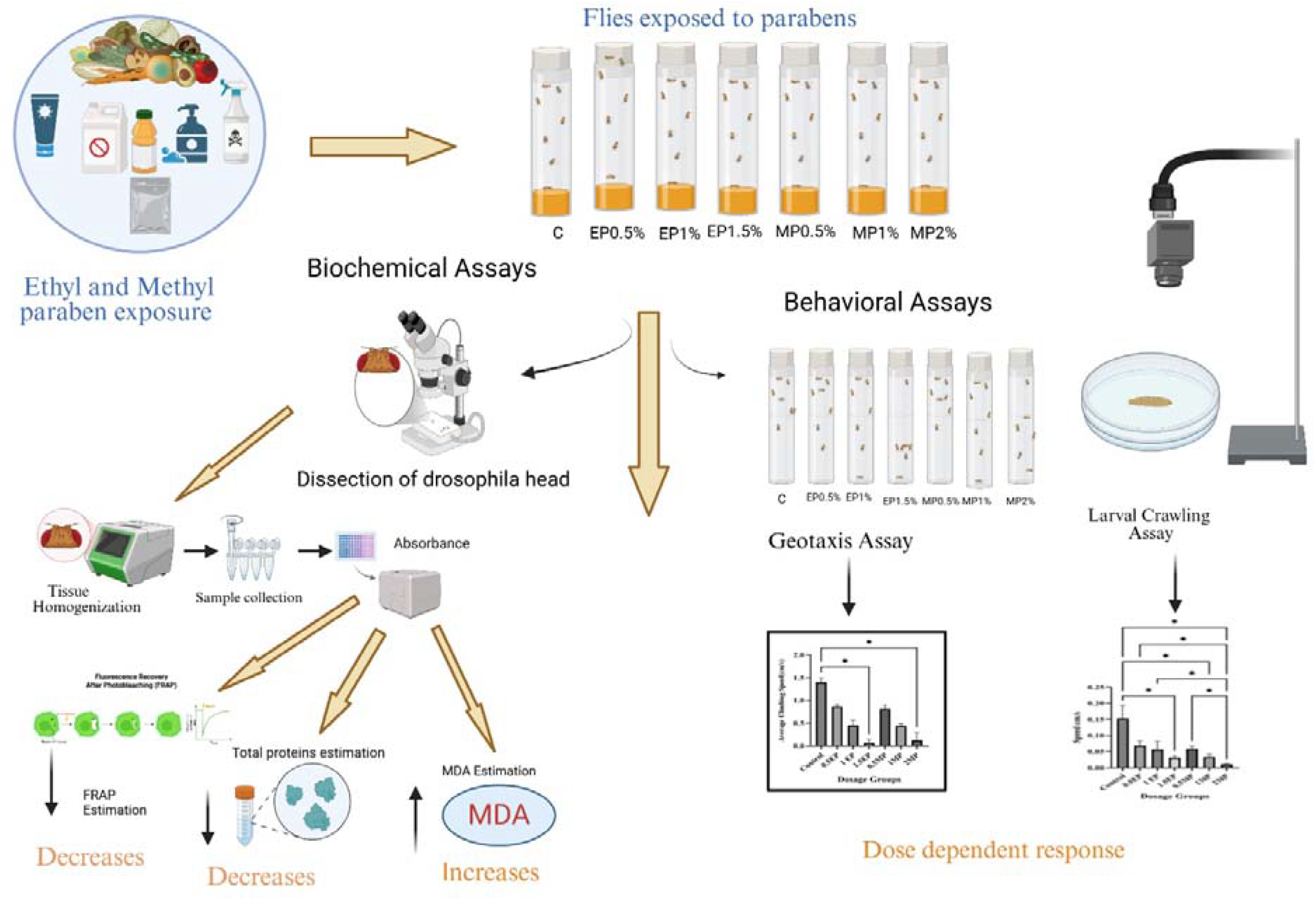
Overview of the experimental design assessing the biochemical and behavioral effects of ethyl- and methyl-paraben exposure in *Drosophila melanogaster*.

**Figure 2A.**
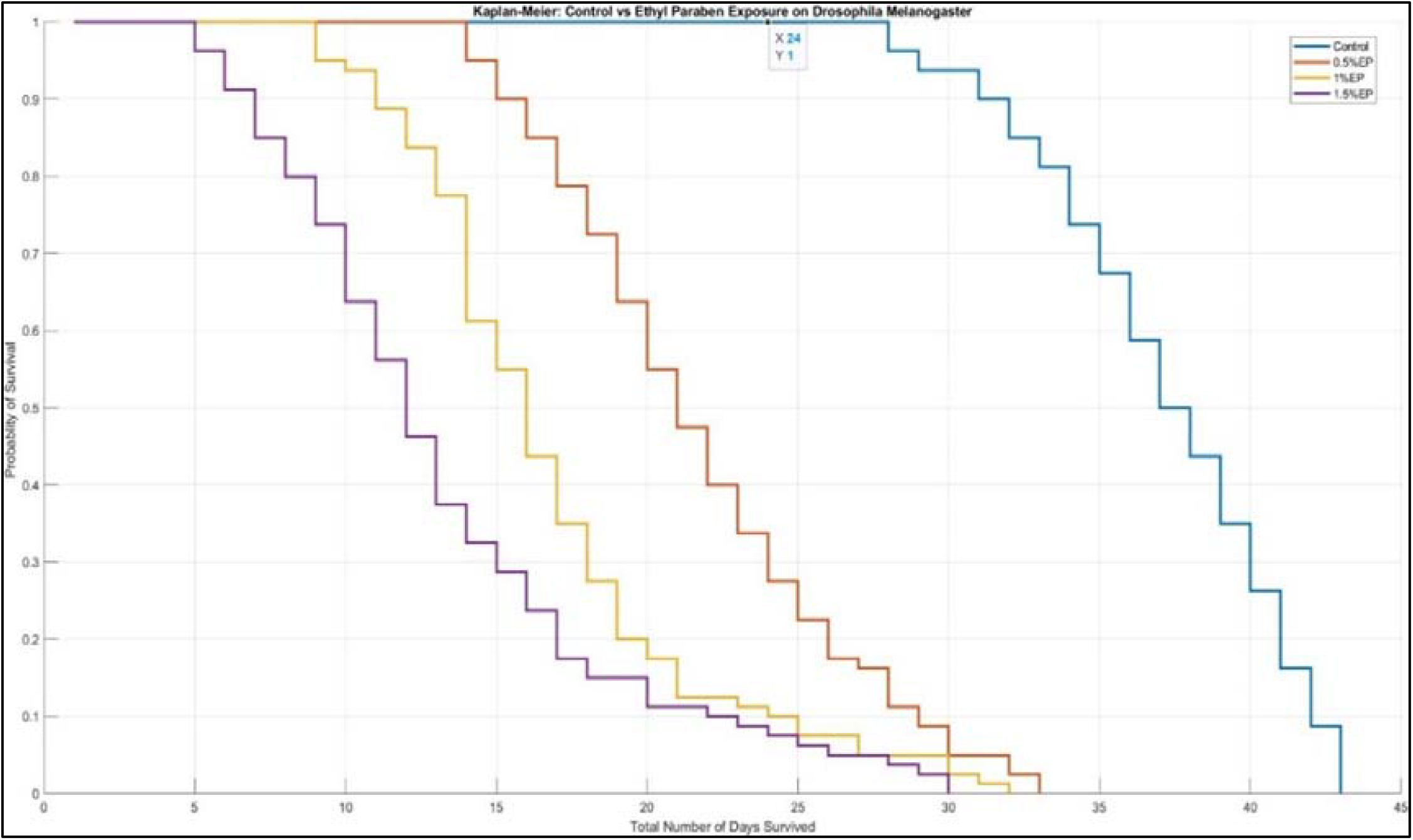
Kaplan-Meier survival curve comparing the effect of ethyl paraben (EP) exposure with the control group on the lifespan of *Drosophila melanogaster*.

**Figure 2b.**
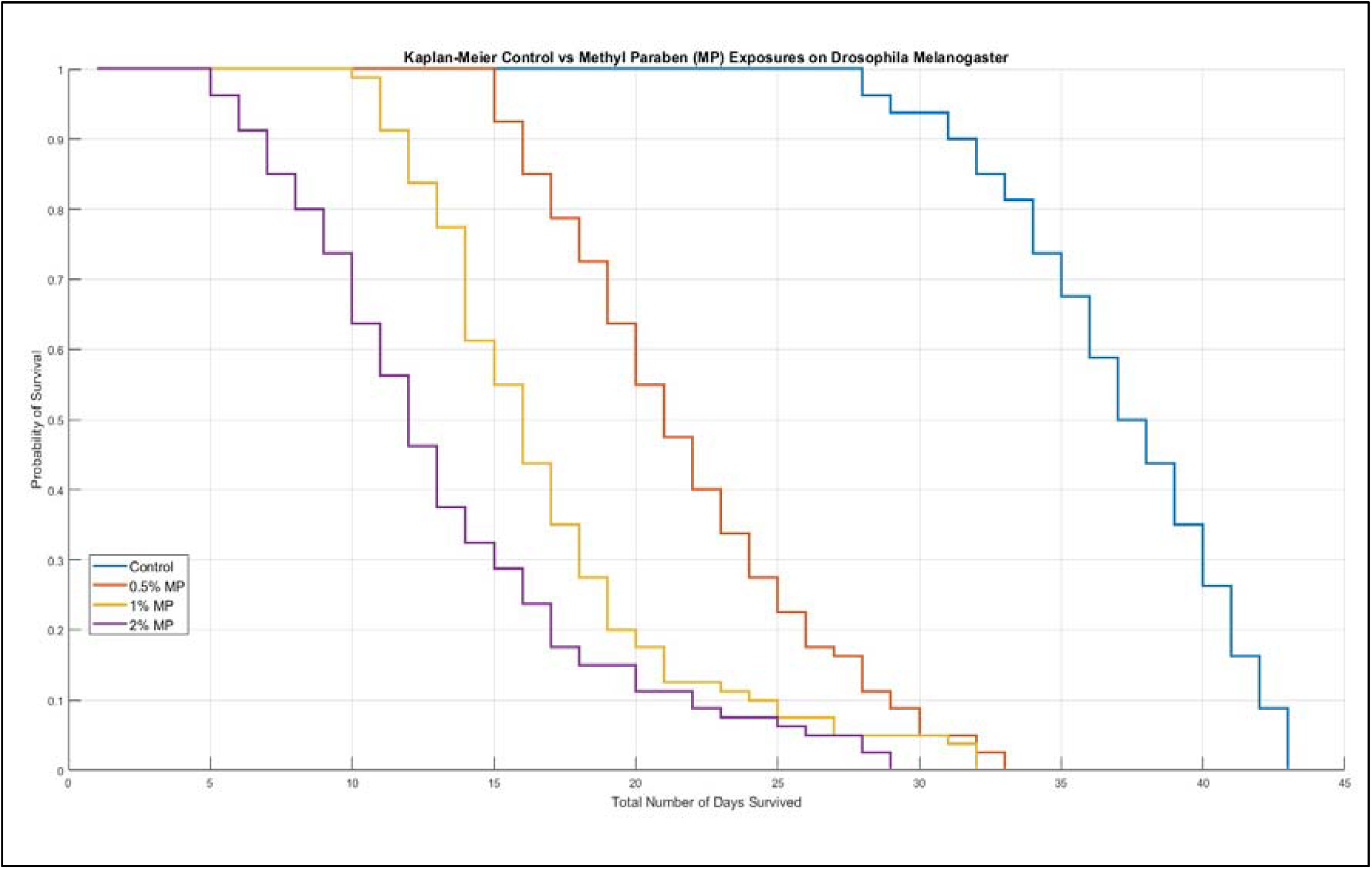
Kaplan-Meier survival curve comparing the effect of methylparaben (MP) exposure with the control group on the lifespan of *Drosophila melanogaster*.

### 4.2. Chronic exposure to paraben induces oxidative damage in flies

Biochemical assays showed that paraben exposure in flies leads to oxidative damage and metabolic disruption. Total protein content in fly heads dropped more rapidly with age in paraben-treated groups, with MP-treated flies (2%) showing about 25% less protein than controls by day 20 (*p*<0.01). This suggests impaired protein synthesis or increased degradation. Flies treated with MP and EP also had *l*ower protein at 30 days. Elevated lipid peroxidation (MDA) and reduced antioxidant capacity (FRAP) were observed in all paraben groups; these effects were dose-dependent (p<0.001 for high doses, p*<0*.*01 for* low doses). MP (2%) caused the highest MDA and greatest FRAP reduction, corresponding to stronger lifespan effects **(Figure 3)**.

**Figure 3.**
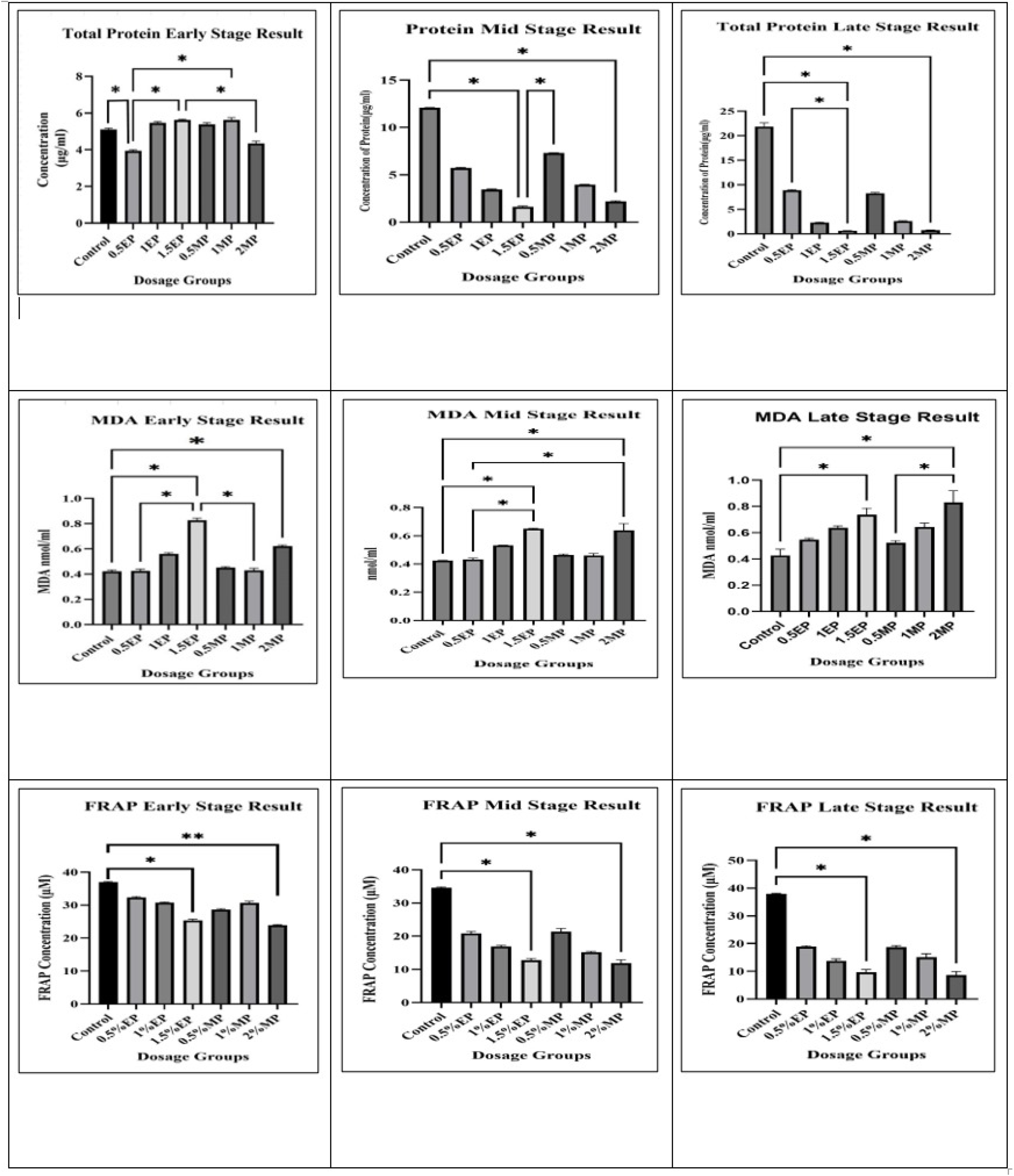
Dose-dependent effects of ethyl- and methyl-paraben exposure on biochemical parameters in *Drosophila melanogaster* across developmental stages.

### 4.3. Paraben Exposure Impacts Locomotory Behavior in Drosophila

Startle-induced negative geotaxis is an innate escape response in adult *Drosophila* that requires intact central nervous system circuits, motor neurons, neuromuscular junctions, and muscle function; as such, it serves as a sensitive, systems-level indicator of neuromuscular integrity. The standardized RING paradigm offers robust, high-throughput, and reproducible measurements of climbing performance and is widely employed to assess age- and stress-related locomotor decline, as well as to investigate genetic or environmental influences. Climbing impairment is a well-established functional hallmark in neurodegeneration models (e.g., α-synuclein overexpression model of Parkinson’s disease flies), highlighting the assay’s translational relevance to motor dysfunction(Gargano et al., 2005). Given that our biochemical endpoints are head-enriched (oxidative-stress markers in neural tissue), coupling them with geotaxis enables the correlation of molecular stress responses to organismal motor behavior within the same cohort.

In the negative geotaxis assay as shown in **Figure 4a and 4b**, control flies demonstrated the highest average climbing speeds during both early and late adult phases (early: 2.35 ± 0.15 cm/s; late: 2.10 ± 0.12 cm/s). Flies exposed to parabens exhibited a dose-dependent reduction in performance. Specifically, early-stage climbing speeds decreased to 2.05 ± 0.13 cm/s (0.5% MP), 1.95 ± 0.14 cm/s (1% MP), and 1.68 ± 0.11 cm/s (2% MP); EP-treated cohorts displayed speeds of **2**.00 ± 0.15 cm/s (0.5% EP), 1.82 ± 0.14 cm/s (1% EP), and 1.60 ± 0.10 cm/s (1.5% EP). This trend persisted into the late stage, with the 2% MP and 1.5% EP groups exhibiting the most substantial deficits (1.50 ± 0.12 cm/s and 1.45 ± 0.10 cm/s, respectively). Kruskal–Wallis analyses revealed significant group differences (early stage: H = 18.97, *p* = 0.00422; late stage: H = 18.70, *p* = 0.00470), while Dunn’s post hoc tests confirmed significant reductions in higher-dose groups relative to controls (*p* < 0.01).

**Figure 4a.**
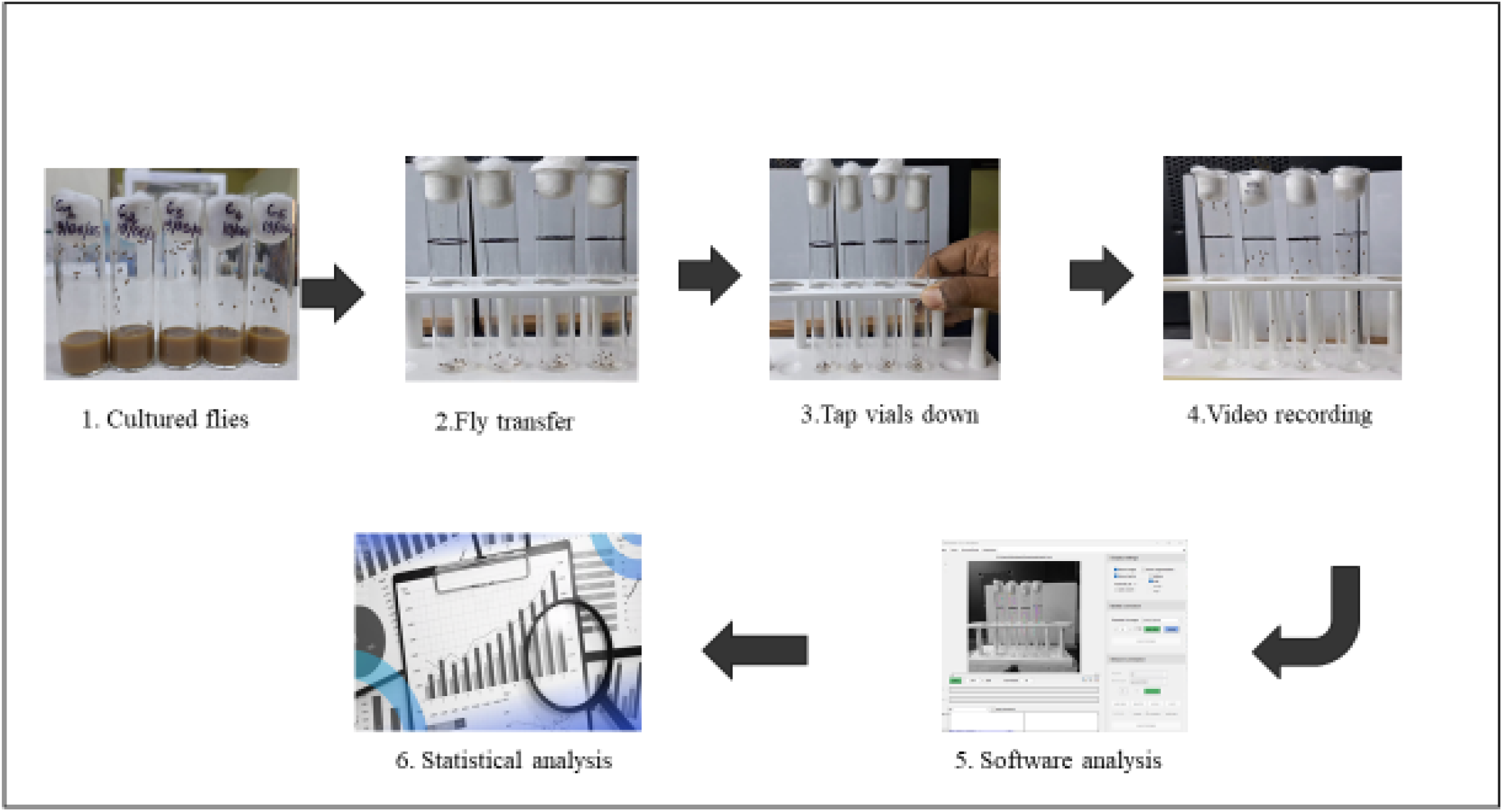
Negative Geotaxis assay procedure.

**Figure 4b.**
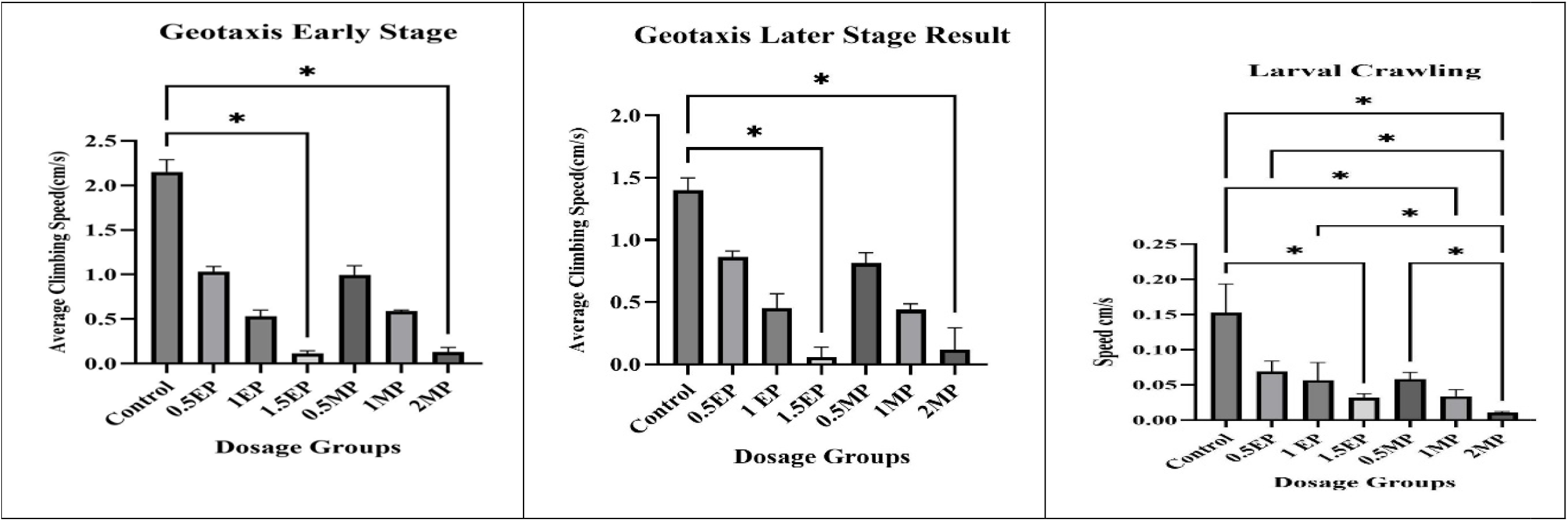
The negative geotaxis assay compares early, mid, and later stages using flies aged 3–5 days, 10–15 days, and 23–25 days, respectively. The larval crawling results demonstrate that both EP and MP influence locomotor activity in a manner dependent on duration and dosage.

To further assess whether locomotor defects manifest earlier developmental stages, we conducted larval locomotion assays **(Figure 5)**. The results indicated that larval crawling assays similarly revealed neuromotor impairment in paraben-exposed specimens. Control larvae had the highest mean crawling speed (3.20 ± 0.18 cm/s), whereas treated groups showed dose-dependent decreases: 2.80 ± 0.15 cm/s (0.5% MP), 2.55 ± 0.14 cm/s (1% MP), 2.30 ± 0.12 cm/s (2% MP), 2.85 ± 0.16 cm/s (0.5% EP), 2.60 ± 0.15 cm/s (1% EP), and 2.35 ± 0.13 cm/s (1.5% EP). The lowest crawling velocity was observed in the 2% MP group, corroborating the greater neurotoxicity associated with higher MP concentrations. Statistical analysis confirmed a pronounced treatment effect (H = 30.10, *p* = 0.000038), with Dunn’s tests showing significant pairwise differences between controls and all higher-dose treatments (*p* < 0.01).

**Figure 5.**
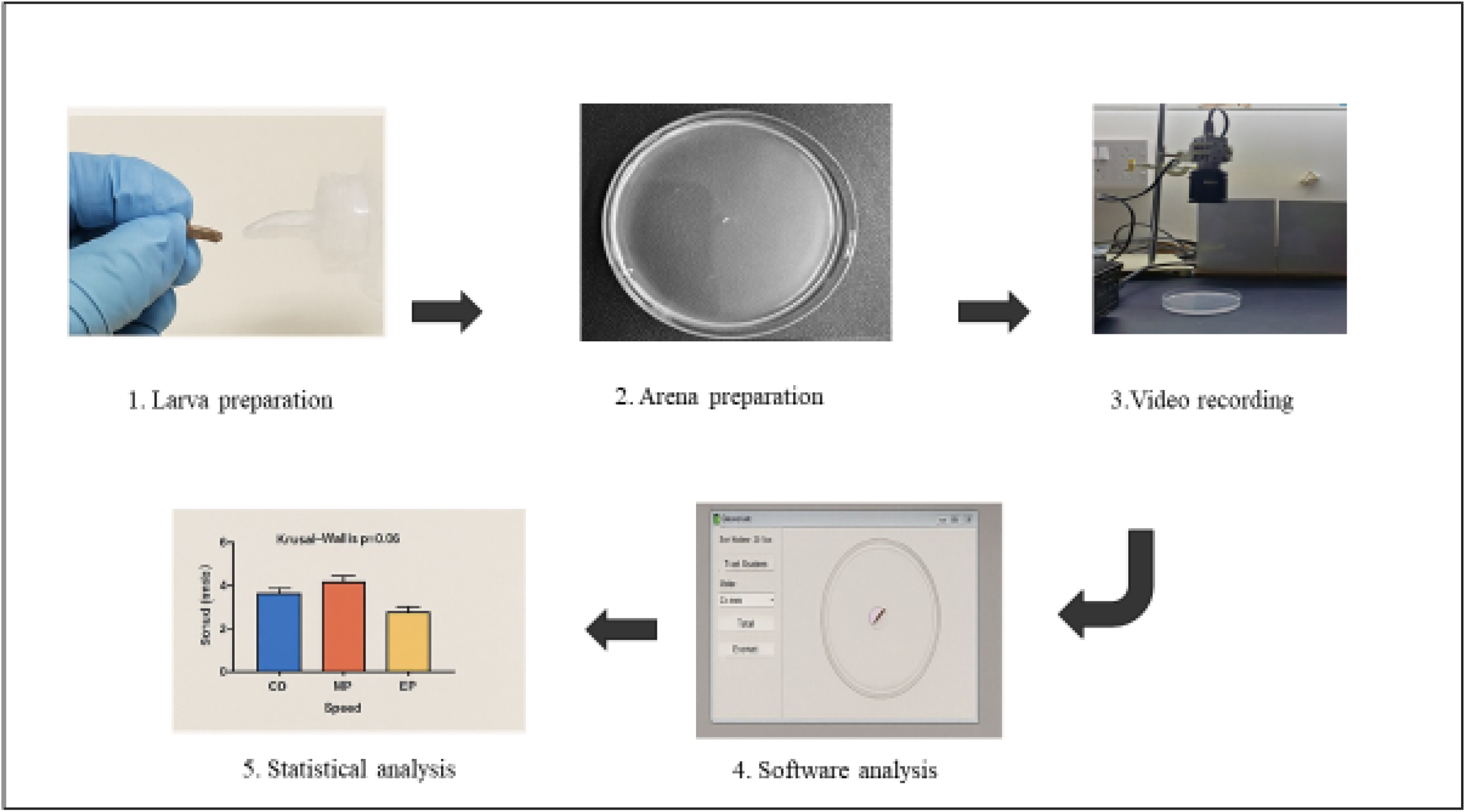
Larval Crawling Assay Procedure.

To eliminate the possibility of nutritional artefacts influencing the results, a feeding assay **(Supplementary Figure 1)** demonstrated equal ingestion of dyed food by both control and treated flies. Consequently, the locomotor impairments detected in both adult and larval stages are attributable to the neurotoxic and neuromuscular effects of paraben exposure, rather than starvation or reduced intake.

## 5. Discussion

### 5.1. Principal findings

Chronic exposure to methylparaben (MP) or ethylparaben (EP) in *Drosophila melanogaster* produced a coherent toxicity profile spanning developmental delay, reduced adult lifespan, head-enriched oxidative stress (↑TBARS/MDA, ↓FRAP), lower soluble protein, and locomotor impairment in both adults (negative geotaxis) and larvae (crawling). Together, these outcomes indicate system-level toxicity that is compatible with endocrine disruption and redox imbalance as upstream drivers.

### 5.2. Endocrine mechanisms

Parabens show weak estrogenic activity, can perturb the hypothalamic–pituitary–thyroid axis, and exhibit anti-androgenic actions in vertebrate models and in vitro, providing credible routes to developmental and reproductive disruption (ERα/ERβ binding; HPT-axis interference; 3α-HSD inhibition)(Kim et al., 2024; Kley et al., 2024). Our developmental delays parallel these pathways and align with prior *Drosophila* data showing that EP can shift ecdysteroid/juvenile-hormone dynamics (delayed/attenuated peaks), a proximate mechanism for slower metamorphosis. However, because we did not quantify ecdysteroids or hormone-responsive gene expression here, endocrine disruption remains an inference rather than a direct demonstration in our cohort. Future experiments should measure 20-hydroxyecdysone titers, EcR/Broad-complex transcripts, and manipulate endocrine nodes genetically to test necessity/sufficiency

### 5.3. Oxidative stress and protein loss

Converging evidence, not direct causation. Across ages (5/20/30 days of exposure) we observed dose-ordered ↑MDA/TBARS and ↓FRAP, consistent with a shift toward pro-oxidant state. In the same heads, soluble protein decreased more steeply with age in treated flies. It is mechanistically plausible that ROS-driven lipid peroxidation co-occurs with protein oxidation/carbonylation and proteolysis, yielding lower recoverable soluble protein; nonetheless, our data is correlative(Dalle-Donne et al., 2003; Niraula & Kim, 2019). Strengthening the causal chain would require assaying protein carbonyls, thiol oxidation, mitochondrial readouts (e.g., aconitase activity, membrane potential), and rescue/epistasis tests (dietary N-acetylcysteine or overexpression of Sod1/Cat) to determine whether dampening ROS normalizes protein content and behavior.

### 5.4. Lifespan effects (dose, structure, and aging biology)

Median lifespan contracted in all exposed groups, with the largest decrement at 2% MP. This pattern is consistent with the well-established contribution of oxidative damage to aging phenotypes and with published evidence that parabens can induce ROS and deplete antioxidants across taxa(Akmal et al., 2024). Still, part of the MP–EP contrast could reflect dose asymmetry (MP tested to 2.0% vs EP to 1.5%) and/or toxicokinetic differences (uptake, metabolism to p-hydroxybenzoic acid). A formal comparison at matched internal doses (e.g., LC-MS/MS of whole-fly paraben/metabolite burdens) would help disentangle structure from exposure.

### 5.5. Neuromuscular/sensory function systems-level readouts

Adult negative geotaxis and larval crawling declined dose-dependently, matching the head-focused biochemical changes. Because geotaxis integrates CNS circuits, motor neurons, NMJs, and muscle, and crawling reflects larval sensorimotor integration, these behavioral assays provide functional linkage between molecular stress and organism-level performance. Paraben-related neurobehavioral effects have been reported in vertebrates (e.g., zebrafish anxiety-like phenotypes)(Clark et al., 2018; Jones & Grotewiel, 2011; Merola et al., 2021), supporting external validity, but mechanisms (estrogenic action vs inflammation/oxidative injury) remain to be parsed. Incorporating electrophysiology (EJPs), synaptic markers, or muscle ultrastructure would sharpen anatomical attribution

### 5.6. Structure–activity consideration

Longer-chain parabens often display stronger estrogenicity/neurotoxicity in mammals; our data suggest that even among shorter chains, MP vs EP differ in magnitude and profile (protein/locomotion vs sensory-leaning effects). Whether this reflects lipophilicity-driven partitioning, membrane penetration, or differential metabolism is testable and relevant to risk assessment, which typically considers parabens as a class(Janiga-MacNelly et al., 2025). Comparative testing at equipotent internal exposures and extension to propyl-/butyl-paraben would position these findings within a Structure activity continuum(Pal et al., 2022).

### 5.7. Translation and exposure context

Humans are widely exposed to parabens through cosmetics, food, and medicines, with measurable levels in blood, urine, breast milk, placenta, etc. Regulatory limits (e.g., ≤0.4% single paraben; ≤0.8% mixtures in cosmetics) were set under assumptions of minimal chronic risk(Hager et al., 2022). Our data show organism-level effects under continuous dietary exposure and strengthen the argument that low-dose, chronic exposures deserve re-evaluation, especially for mixtures and early-life windows(Darbre & Harvey, 2014). While *Drosophila* differs from mammals, many stress and hormonal pathways are conserved, and the fly provides an early-warning platform to triage preservatives and substitutes before costly mammalian studies.

## 6. Limitations

This study only assessed two parabens (MP and EP) at high dietary levels, whereas humans are usually exposed to lower doses and mixtures of parabens. Differences in paraben metabolism between flies and humans could affect dose-responses, as flies lack a liver but have similar detoxification enzymes. Specific hormone levels were not measured to directly confirm endocrine disruption; however, observed developmental delays and gene expression changes strongly suggest this mechanism. Additionally, we focused on continuous exposure from the larval stage, so future work should compare developmental versus adult-only exposures to identify critical periods of susceptibility.

## 7. Implications and Future Directions

Our findings show that parabens are biologically active and affect multiple systems. Using *Drosophila*, we can screen various parabens and substitutes for toxicity. Further studies should examine reproductive effects in *Drosophila*, like mammalian research linking parabens to reduced fertility. Genetic tools could help clarify mechanisms of toxicity by manipulating antioxidant or detoxification genes.

From a public health perspective, our data back calls to reassess paraben safety in consumer products(Commissioner, 2024). Regulatory agencies should factor in cumulative exposure and low-dose effects often missed in standard toxicology tests. In vivo models like *Drosophila* offer fast, ethical insight into chronic impacts. Given widespread paraben use, even small effects may have large population implications. We support reducing exposure and continued research into safer alternatives.

*Drosophila melanogaster* experiments revealed that consumer-level parabens cause oxidative stress, disrupt development, and impair behavior. These results highlight whole-organism risks and reinforce concerns about paraben safety. Consistency with higher organism studies confirms *Drosophila*’s value in toxicology and its usefulness for screening endocrine disruptors. We recommend more integrative studies and a thorough review of exposure guidelines to better safeguard human health.

## 8. Conclusion

Chronic dietary exposure to methyl paraben and ethyl paraben causes significant developmental, biochemical, and behavioral toxicity in *Drosophila melanogaster*. Paraben-treated flies exhibit delayed development and reduced adult lifespan, accompanied by clear signs of oxidative stress (elevated MDA, reduced antioxidant capacity) and protein loss. These biochemical disruptions likely underlie the observed neuromuscular diminished locomotor activity– in exposed flies. Our integrated findings provide mechanistic evidence that links paraben-induced endocrine disruption and oxidative damage to functional deficits in vivo. Given the conservation of many hormonal and stress-response pathways, these results raise concerns that long-term human exposure to parabens could similarly contribute to metabolic and neurobehavioral effects. The study underscores the need to reconsider regulatory safety limits for parabens, considering their endocrine-disrupting potential and cumulative exposure. Furthermore, it highlights the utility of *D. melanogaster* as an effective model for evaluating the systemic impact of environmental chemicals. Continuing to use such models will be important for identifying safer alternatives and protecting human health from subtle yet significant toxicological effects of everyday chemical exposures.

## Acknowledgement

1. Dr. Deepti Trivedi & Staff of Bangalore fly center, Bangalore Fly Centre, NCBS, Bengaluru, Karnataka
2. Dr. Mathivanan Jyothi, Faculty, Department of Human Genetics, NIMHANS, Bengaluru, Karnataka
3. Dr. Sarada S, Head of Department of Neurochemistry, NIMHANS, Bengaluru, Karnataka
4. Dr. Srinivas Bharath MM, Head of Department CPNT, NIMHANS, Bengaluru, Karnataka
5. Dr Farhan Zameer Professor & Chief Scientific Officer (CSO), ATMA Research Centre, Department of Dravyaguna, Alva’s Ayurveda Medical College and Hospital, Moodubidire, India.

## Declaration of Conflict Interests

The authors declared no potential conflicts of interest with respect to the research, authorship, and/or publication of this article.

## Disclosure of Use of Generative AI

None.

## Funding

NIMHANS financial support received for this research.

## Ethical Consideration

IBSC Approval Received.

## Supplemental Material

Supplemental material for this article available online.

## Graphical Abstract

## Notes

### Competing Interest Statement

The authors have declared no competing interest.

