## Supplementary Figure 1 for "Chronic Dietary Exposure to Methylparaben and Ethyl paraben Induces Developmental, Biochemical, and Behavioural Toxicity in Drosophila melanogaster"

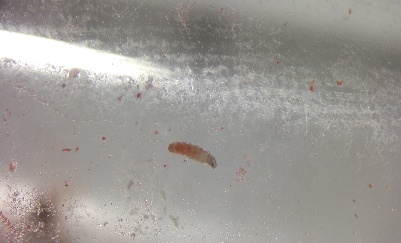

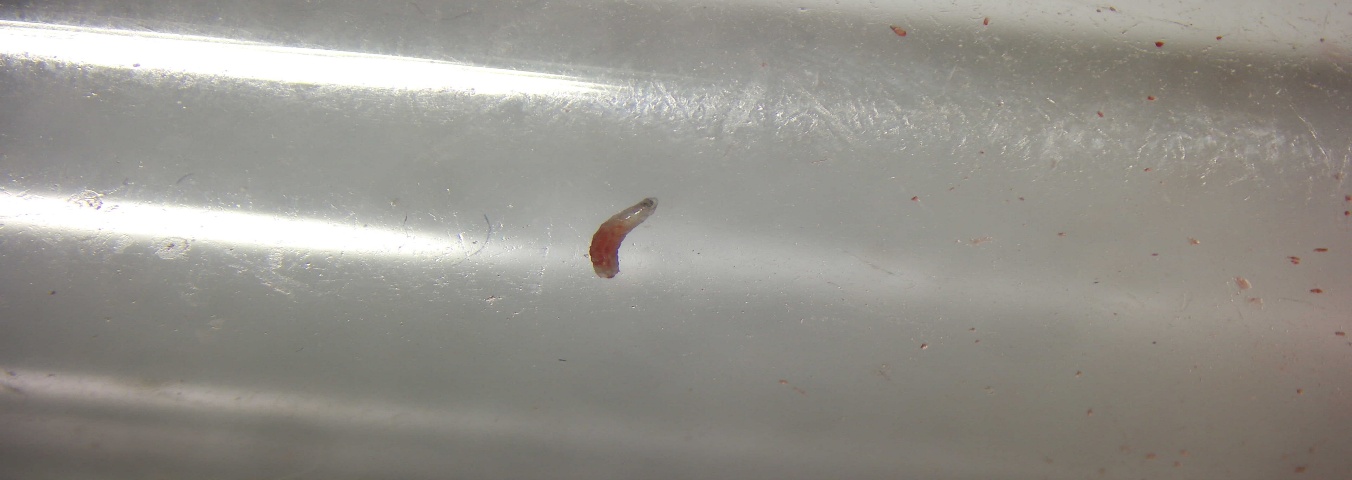

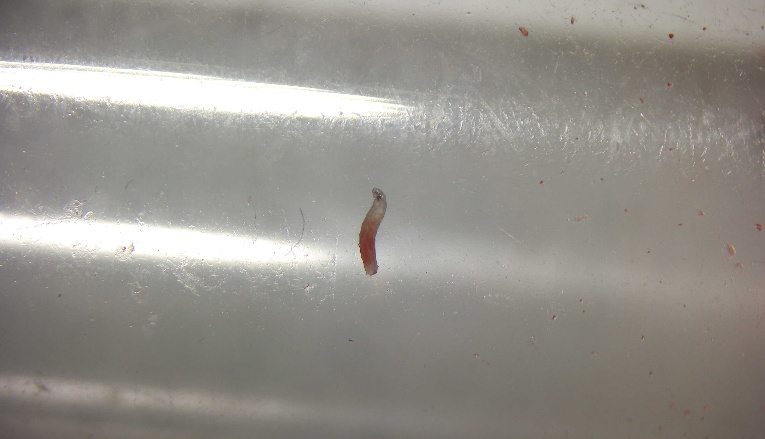

| Geotaxis early stage | Geotaxis later stage | Larval crawling |
| --- | --- | --- |

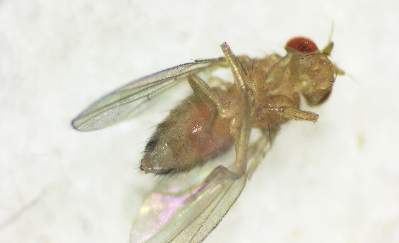

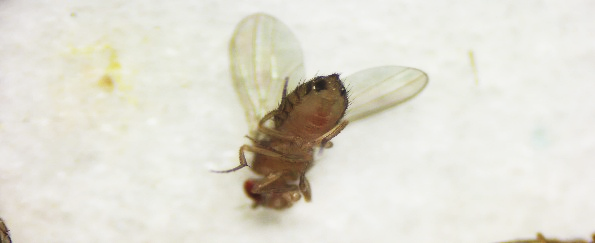

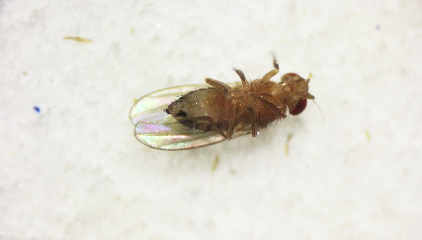

**Supplementary Figure 1** Confirmation of Feeding Assay in *Drosophila melanogaster* Larvae and Adults Across Control and Paraben-Treated Groups Using Red Food Dye.
