## Supplementary Figure 2 for "Chronic Dietary Exposure to Methylparaben and Ethyl paraben Induces Developmental, Biochemical, and Behavioural Toxicity in Drosophila melanogaster"

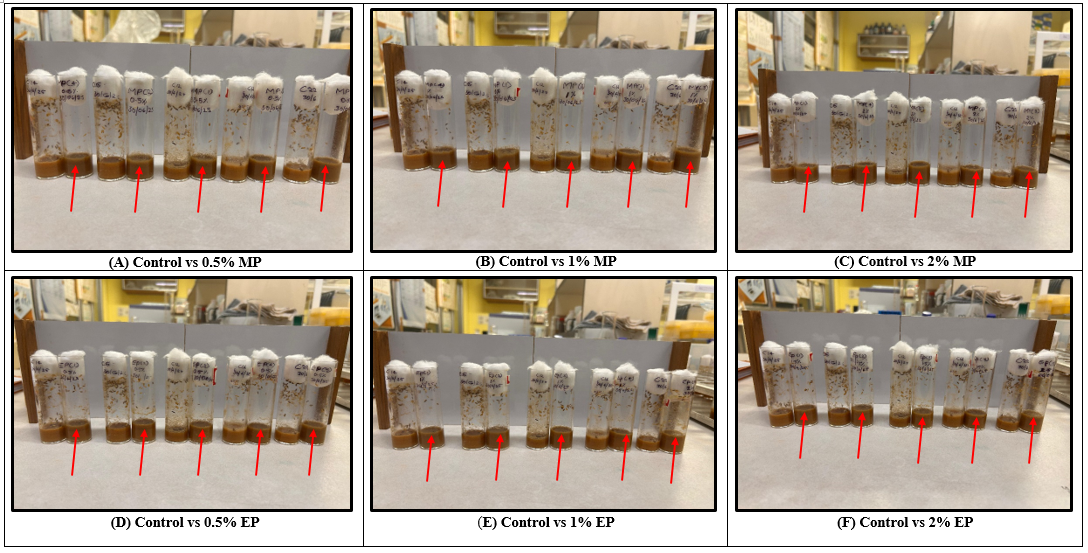
**Supplementary Figure 2**

**Supplementary Figure 2** Visual representation of dose-dependent developmental delay*.* Control groups show normal growth, while increased MP and EP concentrations (panels A–F) result in progressively delayed or arrested development, highlighted by reduced pupal cases and adult flies.
